# Septoria tritici blotch resistance gene *Stb15* encodes a lectin receptor-like kinase

**DOI:** 10.1101/2023.09.11.557217

**Authors:** Amber N. Hafeez, Laetitia Chartrain, Cong Feng, Florence Cambon, Martha Clarke, Simon Griffiths, Sadiye Hayta, Mei Jiang, Beat Keller, Rachel Kirby, Markus C. Kolodziej, Oliver R. Powell, Mark Smedley, Burkhard Steuernagel, Wenfei Xian, Luzie U. Wingen, Shifeng Cheng, Cyrille Saintenac, Brande B. H. Wulff, James K. M. Brown

## Abstract

Septoria tritici blotch (STB), caused by the Dothideomycete fungus *Zymoseptoria tritici*, is of one of the most damaging diseases of bread wheat (*Triticum aestivum*)^1^ and the target of costly fungicide applications^2^. In line with the fungus’ apoplastic lifestyle, STB resistance genes isolated to date encode receptor-like kinases (RLKs) including a wall-associated kinase (*Stb6*) and a cysteine-rich kinase (*Stb16q*)^3,4^. Here, we used genome-wide association studies (GWAS) on a panel of 300 whole-genome shotgun-sequenced diverse wheat landraces (WatSeq consortium) to identify a 99 kb region containing six candidates for the *Stb15* resistance gene. Mutagenesis and transgenesis confirmed a gene encoding an intronless G-type lectin RLK (LecRK) as *Stb15*. The characterisation of *Stb15* exemplifies the unexpected diversity of RLKs conferring *Z. tritici* resistance in wheat.

## Main

The domestication of wheat 10,000 years ago heralded the dawn of modern agriculture in western Eurasia^5^ whilst providing an opportunity for the specialisation of an uninvited guest: the fungal pathogen *Zymoseptoria tritici*^6^. Understanding and bolstering genetic resistance could aid in reclaiming ∼24 million tonnes of yield lost to STB each year^1,7^.

During its interaction with wheat, *Z. tritici* colonises the apoplast through the stomata and commences a period of asymptomatic growth wherein effectors are released: molecules that suppress host defences or make the host amenable to colonisation^8^. Host resistance proteins may directly or indirectly recognise these effectors and modulate defence responses, described in apoplastic interactions as effector-triggered defence or the ‘invasion model’^9–11^. If undetected, the pathogen switches to its necrotrophic life stage, resulting in the release of host nutrients and the rapid growth and proliferation of the pathogen^12^. Symptoms ultimately manifest as necrotic lesions on the leaf surface containing pycnidia (asexual fruiting bodies), which produce conidia that may disperse up to one metre by rain splash, allowing further cycles of colonisation and thus quick progress of the disease^13^.

Twenty-three major genes controlling isolate-specific resistance to STB (*Stb* genes) have been mapped in wheat^14–16^, but *Stb* gene cloning has lagged behind efforts for other wheat diseases. *Stb6* on chromosome 3AS, conferring race-specific resistance to *Z. tritici*^3,17^ encodes a wall-associated receptor kinase (WAK), a subfamily within the receptor-like kinase (RLK) family in plants, with a galacturonan-binding domain^3^. *Stb16q* on chromosome 3D^18^ encodes a cysteine-rich receptor kinase (CRK) with two DUF26 domains^4^. Thus, the two *Stb* genes cloned to date encode RLKs with extracellular domains which have a putative sugar-binding function. *Stb15* is a major gene for resistance to *Z. tritici* isolate IPO88004, mapped to a 36 cM region in the cultivar Arina^19^. It is a good candidate for cloning due to its large phenotypic effect resulting in full resistance, which is rare amongst *Stb* genes^14^, and is important due to its presence across the breadth of European wheat cultivars^20^.

Here we apply genome-wide association studies (GWAS) to map resistance to STB in the diverse Watkins collection of pre-Green Revolution wheat landraces, which provides the opportunity to study interactions with STB in a well-adapted yet highly genetically diverse context^21,22^. GWAS harnesses naturally-occurring population structures in collections of accessions representing the genetic and phenotypic diversity of a species^7,23^. Well-curated and sequence-configured panels can be tested for correlations with multiple phenotype datasets to potentially map many genes from a single population. For a truly unbiased approach, whole-genome shotgun (WGS) sequencing can be employed to access the entire genetic diversity of a panel. Sequence reads can then be aligned to a reference genome and the resulting SNP calls used for GWAS.

STB symptoms elicited by the *Z. tritici* isolates IPO323, avirulent to *Stb6*^17^, and IPO88004, avirulent to *Stb15*^19^, were scored across 300 Watkins landraces (**Fig. 1a**). This core panel was selected to maximize genetic representation^24^. Leaf damage (necrosis and chlorosis), and pycnidial coverage are usually, but not always, correlated^25^. Both phenotypes were recorded at 5-6 timepoints for calculation of the area under the disease progress curve (AUDPC), followed by logit transformation and linear mixed modelling (**Supp. Tables 1-4**).

**Fig. 1:**
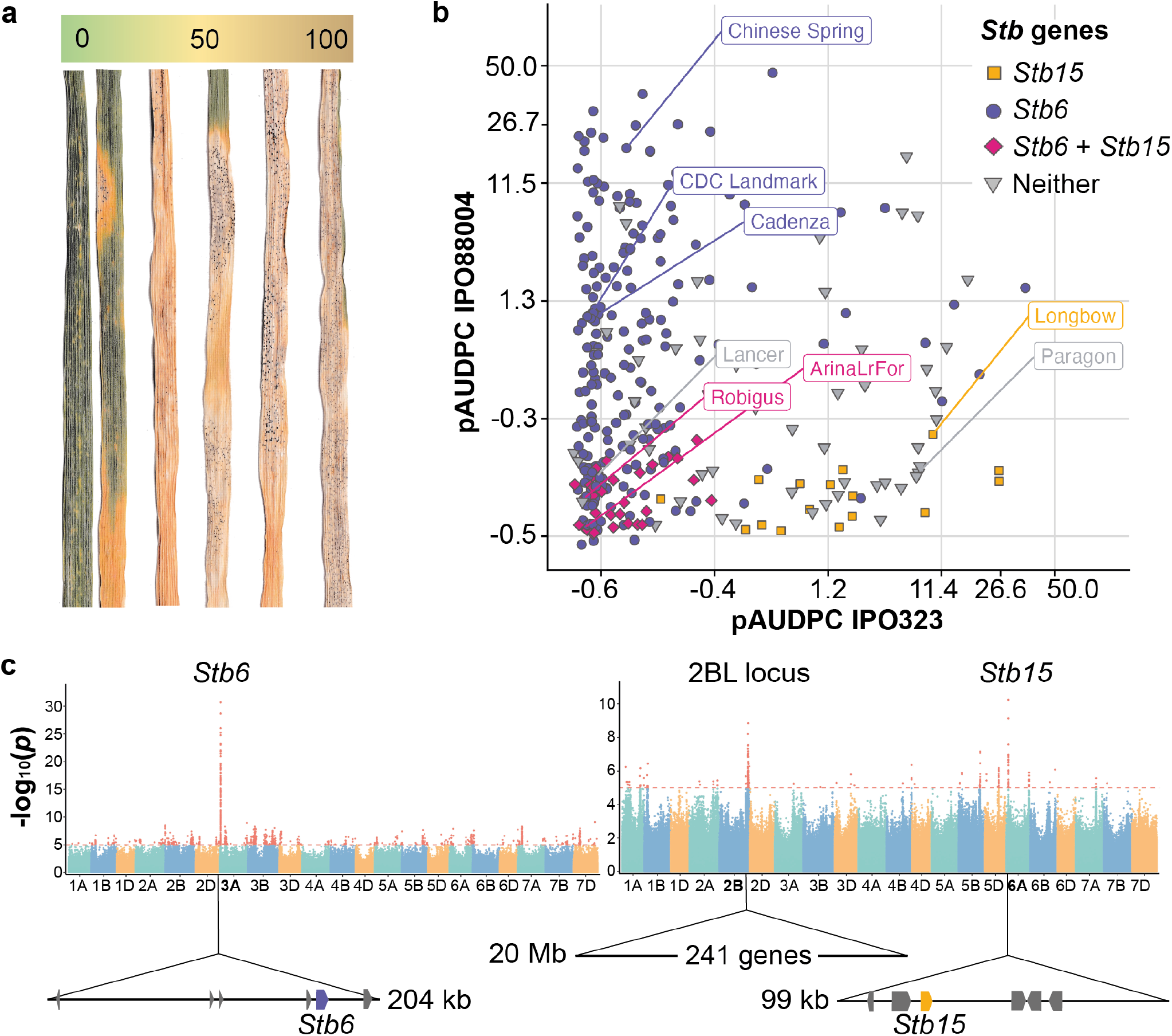
Race-specific resistance to *Z. tritici* in the wheat Watkins landrace panel associates with discrete disequilibrium blocks. **a**, Quantitative variation in pycnidia and necrosis phenotypes. Pictured are leaves arranged by pycnidia coverage. **b**, Effects of *Stb6* and *Stb15* on resistance to *Z. tritici* isolates IPO88004 and IPO323. **c**, Manhattan plots showing the association of logit pAUDPC response to *Z. tritici* isolates IPO323 (left) and IPO88004 (right) with SNPs mapped to Chinese Spring. Linkage disequilibrium (LD) blocks associated with STB resistance are drawn as arrows beneath the chromosomes (marked in bold) with the 6A *Stb15* candidate gene marked in orange and *Stb6* marked in purple.

A SNP matrix generated from WGS sequencing data of wheat cultivars and landraces was mapped to Chinese Spring and employed for GWAS (WatSeq consortium). As a positive control to ensure the suitability of the experimental system for mapping via this method, *Stb6* was successfully restricted to a discrete genomic interval in the core Watkins panel. We screened IPO323 on the Watkins core panel and ten control cultivars (**Supp. Table 5**), including Chinese Spring which has the functional allele of *Stb6*^3^ (**Fig. 1b**). An interval on chromosome 3A was associated with both leaf damage and pycnidia phenotypes (pycnidia: **Fig. 1c**; leaf damage: **Supp. Fig. 1**). SNPs in the 3A locus were highly associated with pycnidia, with a -log10 *p*-value of almost 30. Within this region, a linkage disequilibrium (LD) block extending from 26.10 to 27.50 Mb was identified. A smaller haploblock within it was most highly associated with resistance, from 26,035,170 to 26,238,727 bp (**Supp. Fig. 2**). This 203.6 kb region contained six genes, including *Stb6* (**Fig. 1c**).

We then proceeded to identify *Stb15* by inoculating the panel with IPO88004 and employing GWAS. Several regions were associated with STB phenotypes (pycnidia: **Fig. 1c;** leaf damage: **Supp. Fig. 3**). A locus on 6AS had the highest *p*-value for both pycnidia and damage traits and spanned a 99.1 kb region between 485,503,26 and 485,994,21 bp containing six genes. Comparison of gene sequences between Arina*LrFor* and Chinese Spring combined with correlation of haplotypes with the responses of landraces to IPO88004 excluded five of these genes (**Supp. Table 6**; **Supp. Fig. 4**). The remaining gene, TraesCS6A02G078700/TraesARI6A03G03215890, is predicted to encode an RLK and is strongly associated with isolate-specific resistance to IPO88004 (**Fig. 1b**), so was selected as the most likely candidate for *Stb15*.

We also observed a significant association of pycnidia cover of IPO88004 with a locus on chromosome 2BL which spanned 755 to 775 Mb and contained 241 genes. When we removed the masking effect of lines carrying *Stb15*, the significance of the 2BL resistance increased one-thousand-fold (**Supp. Fig. 5**). *Stb9* has previously been mapped to 2BL^26^ but is outside of this locus (at ∼808 Mb)^27^ and accessions which display resistance to IPO89011, an isolate which is avirulent on *Stb9*, are not always resistant to IPO88004^20,28^ (**Supp. Table 7**). Therefore, the LD block appears to be a novel locus for resistance to *Z. tritici*, temporarily designated as *STBWat1*.

TraesCS6A02G078700/TraesARI6A03G03215890 was confirmed as *Stb15* by a combination of mutagenesis and transgenesis and shown to be a lectin receptor kinase (LecRK). We screened 3,308 plants from 307 M2 families of an EMS-derived mutant population of cv. Arina for resistance to IPO88004 and identified three independent susceptible mutants (**Fig. 2a,b**). All three of these mutant plants had one non-synonymous transition mutation in the open reading frame of the *Stb15* candidate. The gene encodes a G-type lectin receptor kinase (LecRK)^29^ with an intracellular serine/threonine receptor-like protein kinase (S/TPK) and three extracellular domains: a mannose-specific bulb-type lectin (BTL), an S-locus glycoprotein (SLG), and a plasminogen/apple/nematode (PAN) domain. All three of the induced mutations resulted in replacement by larger amino acids in the BTL and kinase domains. In an AlphaFold model (**Fig. 2c**), all three residues were in locations where mutations would be predicted to cause disruption to the protein structure. To confirm the function of the candidate gene we synthesized a 10.9 kb genomic sequence containing 2 kb and 1.5 kb of 5’ and 3’ regulatory sequence from Arina into a binary vector and transformed wheat cv. Fielder, which is susceptible to isolate IPO88004. We obtained two independent homozygous single-copy T_2_ transgenic lines which conferred resistance to IPO88004 whereas their respective nulls were susceptible, indicating that the isolated gene sequence is sufficient to confer the *Stb15* phenotype (**Fig. 2d; Supp. Table 8-9**). Transgenic lines with four and six to eight copies of *Stb15* were also resistant relative to the controls.

**Fig. 2:**
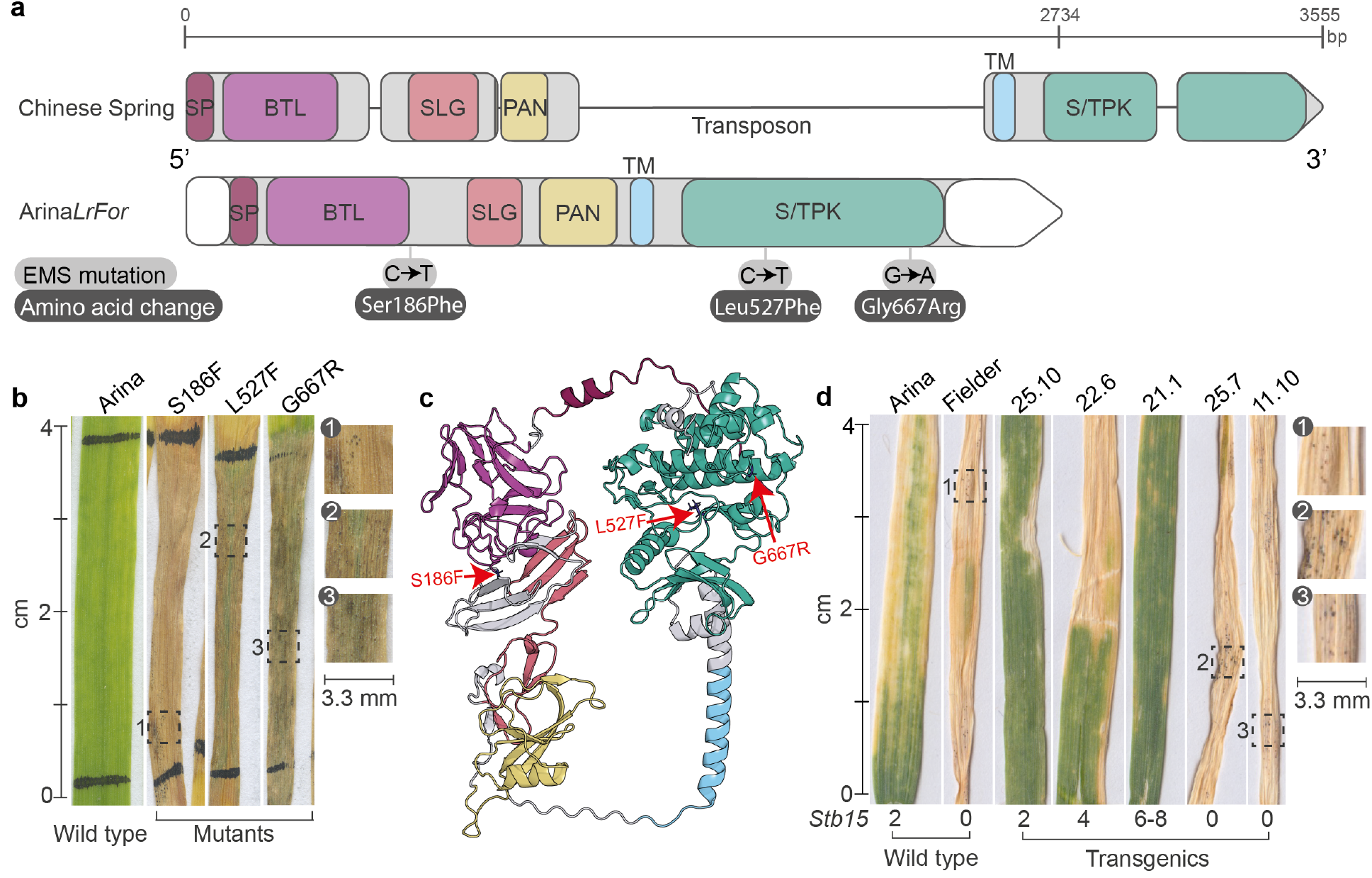
Structure and function of *Stb15*. **a, b, a**,**b** The functional resistance allele of *Stb15* in wheat cv. Arina and Arina*LrFor* compared to the susceptible allele in cv. Chinese Spring. The predicted exons and introns are shown as rounded rectangles and lines, respectively, for Chinese Spring (RefSeq v1.1^75^) and Arina*LrFor* (see Methods). Domains are highlighted: SP = signal peptide, BTL = bulb-type lectin, SLG = S-locus glycoprotein, PAN = plasminogen/apple/nematode, TM = transmembrane, S/TPK = Serine/Threonine Protein Kinase. White boxes indicate untranslated regions (UTRs). The sequence and phenotype of three EMS-induced loss-of-function mutants inoculated with *Z. tritici* isolate IPO88004 are indicated. **c**, AlphaFold-augmented 3D structural model of Stb15. The domains are coloured as in panel a. The location of the three EMS-induced mutations are indicated by dark blue colouring and red arrows with labels. **d**, Cultivar Fielder stably transformed with an *Stb15* construct and inoculated with isolate IPO88004. 25.7 is a null wherein the transgene segregated out in the T_2_ family whilst 11.10 was transformed with the same vector backbone minus *Stb15*. Copy number of *Stb15* (*Stb15*) is given as a fixed number or range.

The functional allele of *Stb15* is present across the geographic (**Fig. 3a**) and genetic (**Fig. 3b**) diversity of the Watkins core 300 collection, although it occurs in only 15% of landraces. It is often present alongside *Stb6*, which is more common (78%). Alleles were defined based on SNP distance (**Supp. Fig. 4**; **Supp. Table 10-11**). 14% of landraces displayed resistance to *Z. tritici* which could not be explained by either gene. Unexplained resistance to IPO88004 (36 landraces) could be due to *STBWat1* (**Fig. 1c**). *Stb15* is also present in 35% of European cultivars tested using KASP markers (**Supp. Table 12**).

**Fig. 3:**
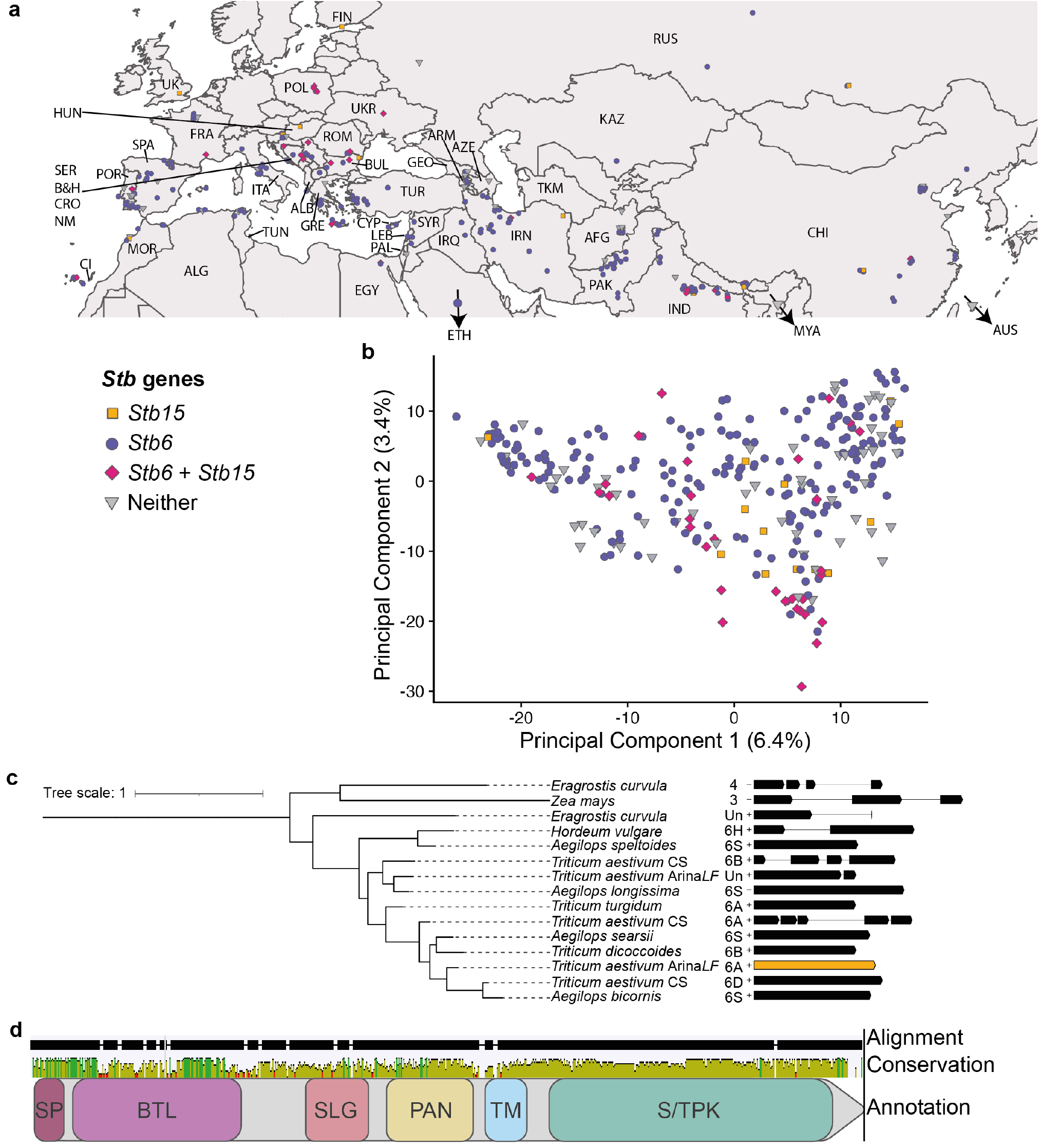
Geographic distribution and intra- and inter-species structural diversity of *Stb15*. **a**, Distribution of *Stb6* and *Stb15* in the Watkins 300 core collection. The map indicates coordinates of markets from which landraces were collected. Only countries from which landraces were collected are labelled. Country abbreviations are expanded in **Supp. Text 1. b**, Principal Component Analysis (PCA) plot of 300 Watkins landraces with lines containing predicted functional alleles of *Stb6* (purple), *Stb15* (orange), both (pink) or neither (grey) indicated. **c**, Maximum likelihood phylogenetic tree of proteins with homology to the Arina*LrFor* (Arina*LF*) Stb15-encoded allele from selected Poaceae species, including the wheat reference genome Chinese Spring (CS). The smallest non-repetitive (‘inner’) clade containing *Stb15* is shown. The intron/exon structure of *Stb15* homologs and their relative nucleotide lengths are presented (arrow = exon coding sequence, line = intron). Species names and chromosomes are given; Un indicates homologs within scaffolds which have not yet been mapped to chromosomes. Gene IDs for homologs are given in **Supp. Table 14. d**, Protein alignment of the homologs from **panel c** with alignment gaps, sequence conservation and predicted protein domains indicated. Taller, greener bars in the conservation panel indicate more conserved regions.

Forty-eight proteins from 16 Poaceae species shared homology with the Arina/Arina*LrFor* Stb15 protein (**Supp. Fig. 6**; **Supp. Table 13)**. Homologous genes encoding the protein sequences were found to be conserved across the Group 6 chromosomes within the Triticeae (**Fig. 3c**), but were also present on other chromosomes, especially Groups 3, 4 and 7 (**Supp. Fig. 6**).

We detected intron/exon structural diversity in gene annotations of proteins with homology to Stb15 across the Poaceae, including within *T. aestivum* (**Fig. 3c**). The functional allele of *Stb15* is intronless, whilst in cv. Chinese Spring, lacking the *Stb15* phenotype, the gene has four introns. An intronless gene structure was also observed in gene annotations of 22 homologous proteins both within the Triticeae tribe (*Triticum, Aegilops* and *Hordeum*) and beyond it (*Brachypodium* and *Avena*), especially on Group 3 and 6D/6S chromosomes. *Stb15* clustered most closely to the Chinese Spring 6D and *Aegilops bicornis* 6S homologs, suggesting that the functional allele of *Stb15* may have originated from the D or S genomes, which share high sequence homology^30^.

Within the inner *Stb15* clade, the kinase and PAN domains were highly conserved whilst the region spanning the BTL and SLG domains was variable (**Fig. 3d**), suggesting it may be under diversifying selection.

Characterisation of the third *Stb* gene from a distinct subclass of the RLK protein family has the potential to enhance molecular understanding of the wheat-*Z. tritici* interaction, providing new opportunities for research and disease control. In addition, this research demonstrates the power of GWAS to greatly accelerate gene cloning for traits which are poorly understood at the molecular level. One of the factors that may limit the success of GWAS is population structure^31^. In this study, the presence of *Stb15* and *Stb6* in Watkins landraces that spanned the breadth of the genetic diversity of the panel was likely decisive to their successful mapping (**Fig. 3a,b**). Such wide distributions of allelic variation across the full range of relevant germplasm allow the effects of genes of interest to be separated from those of kinship. Such a distribution may be more likely for genes which were introduced early into cultivated hexaploid wheat, which appears to be the case of *Stb6* known from both Europe and East Asia^32^. Likewise, Watkins lines with *Stb15* were obtained across the breadth of Eurasia as well as North Africa.

The diversity of intron/exon gene structures amongst *Stb15* homologs is unusual when compared to nucleotide-binding leucine-rich repeats (NLRs) for which gene structures tend to be conserved^33^. Leucine-rich repeat membrane-anchored proteins without intracellular kinases control resistance to the related Dothidiomycete fungus *Cladosporium fulvum*, the causal agent of leaf mold in tomato^34^, many of which^35^ share the intronless open reading frame exhibited by the functional allele of *Stb15*. Possibly, intronless gene structures have been conserved whilst intron gain has occurred in *e*.*g*. the Chinese Spring allele.

*Stb15* has more extracellular domains compared to *Stb6* and *Stb16q*, and the diversity of RLK subclasses conferring resistance is unusual compared to *Cf* genes but similar to genes conferring resistance to blackleg disease in *Brassica* spp.^36,37^. Equally, there are similarities shared by Stb proteins: they are transmembrane proteins with extracellular domains with a putative sugar-binding function and an intracellular kinase. Both the DUF26 domains of *Stb16q* and the G-type lectin domain of *Stb15* likely bind mannose, a building block of mannan found in cell walls of both fungi^38^ and wheat^39^. Detection of a conserved PAMP fits the function of lectins which form part of basal plant immunity and are involved in stomatal innate immunity responses in *Arabidopsis thaliana*^40^, and LecRKs confer non-host or marginal host resistance to leaf rust in barley^41^. This role could make Stb15 a target for suppression by the pathogen or it could be part of a guard/guardee pair^42^, triggering isolate-specific resistance. Alternatively, the interaction could resemble that of the tomato receptor Cf-4 and Avr4, a passive *C. fulvum* effector which binds chitin to avoid breakdown by the host plant^34^. Another possibility is that *Stb15* binds glycoproteins; AFP1 in maize was previously thought to bind chitin but in fact interacts with chitin deacetylases, most likely via their mannosylated group^43^.

LecRKs have been found to bind to secreted proteins, *e*.*g*. a *Phytophthora spp*. effector^44^, which may also be the case for Stb15. A candidate gene for *AvrStb15* encoding a small secreted protein (SSP) has been suggested^45^, but further work will be needed to determine the nature of its interaction with Stb15. There is thus far no evidence of a direct interaction between Stb6 and AvrStb6, also encoding a cysteine-rich SSP^3,46,47^. In conclusion, our study highlights the importance of elucidating the diverse roles of Stb-AvrStb pairs in defence induction for understanding the genetic basis of resistance in this economically important pathosystem.

## Acknowledgements

We thank Cristobal Uauy, Anthony Hall and Manuel Spannagl for pre-publication access to Arina*LrFor* RNAseq data and associated genome annotations, Tom O’Hara for sharing resources, Tjelvar Olsson for Python training and consultation and Sreya Ghosh and Guru Radhakrishnan for advice on phylogenetics.

This research was supported by the NBI Research Computing group, the Informatics Platform and Horticultural Services at the John Innes Centre, UK, the experimental infrastructure VégéPôle, INRAE, France, and financed by funding from the Biotechnology and Biological Sciences Research Council (BBSRC) Plant Health (BBS/E/J/000PR9798) and (BBS/E/J/000PR9780) Designing Future Wheat Strategic Programmes to J.K.M.B. and B.B.H.W., King Abdullah University of Science and Technology to B.B.H.W and the BBSRC Doctoral Training Program (DTP) at the Norwich Research Park (NRP) to A.N.H.

## Draft of author contributions

Authors are listed in alphabetical order by last name apart from the first three and last four authors. *Z. tritici* pathology work on Watkins landraces was planned and implemented by A.N.H, L.C. and R.K. with supervision from J.K.M.B. Statistical analysis of pathology work was carried out by A.N.H and J.K.M.B. Pre-publication collaboration (within ‘WatSeq’), access to genotyping data and application of GWAS to the project was facilitated primarily by S.C. alongside S.G. and implemented by L.U.W., C.F., W.X. and M.J., as well as generation of associated figures and suggestion of candidate genes. A.N.H and B.S. analysed WatSeq genotypes for allele characterisation and identification of candidate genes. Cloning strategy was conceived and planned by B.B.H.W. An Arina EMS mutant population was generated by M.C.K. in the labs of B.K. Large-scale screens were planned and implemented by C.S. and F.C, along with identification and resequencing of induced susceptible mutants and provision of phenotype images. O.R.P. generated and annotated the AlphaFold model of *Stb15*. M.S. designed a vector carrying *Stb15* which was transformed into wheat by S.H. and M.C., followed by SSD to T_2_ and screening with *Z. tritici* by L.C. and A.N.H. and statistical analysis by J.K.M.B. Study of *Stb15* homologs and gene structural variation was conducted by A.N.H. Composite figures were designed by A.N.H., B.B.H.W. and J.K.M.B. and generated by A.N.H. A.N.H drafted the manuscript with extensive input and revisions from B.B.H.W., J.K.M.B. and C.S. Further revisions to the manuscript were contributed by B.K., M.C.K., L.C., S.H., M.S., C.F. and M.J.

## Materials and Methods

### Plant and pathogen material

Of the total 826 lines in the Watkins collection, we used a core set of 300 lines representing the majority of genetic variation present in spring growth types^24^. Wheat control lines were included in all assays, with lines selected based on known response to Septoria or strategic importance (**Supp. Table 5**). Both Arina and Arina*LrFor* were used for analyses or experiments pertaining to *Stb15* as they carry the same allele. Arina*LrFor* is a genotype derived from a cv. Arina x cv. Forno cross and further backcrossing with Arina^48^, where cv. Forno is susceptible to IPO88004, so the resistance in ArinaLrFor should come from the Arina allele of *Stb15*. Seeds of wheat cultivar Arina*LrFor* (PANG0001) are available from the Germplasm Resources Unit, John Innes Centre, Norwich, UK (https://www.jic.ac.uk/research-impact/germplasm-resource-unit/). The *Z. tritici* isolates IPO323 (virulent on *Stb6*) and IPO88004 (virulent on *Stb15*) were used due to known avirulence to *Stb6*^3,17^ and *Stb15*^19^, respectively. IPO323 was isolated in 1981 in The Netherlands^49^ whilst IPO88004 was isolated in Ethiopia in 1988^50^. A third isolate, IPO90012 from Mexico^51^, was also included for comparison as a virulent control isolate.

### Design and infection protocol for pathology assays at JIC

An alpha lattice design was used for the pathology experiments which consisted of five replicates across incomplete blocks (40-well seedling trays). This allowed the effects of tray and position in the controlled environment room (CER) to be estimated through statistical analysis. The design was generated using the ALPHA setting of the Gendex programme (http://designcomputing.net/gendex/) based on the design principles of Patterson and Williams (1976)^52^.

The following methods are based on those described by Arraiano et al. (2001)^53^, which in turn followed the methods of Kema et al.^50^. Multiple seeds of the lines tested were pre-germinated in Petri dishes on filter paper (Whatman 90 mm, Whatman International Ltd, Hadstone, UK) containing 4 ml of 0.2 ppm gibberellic acid. Petri dishes were placed in the dark at room temperature for 48 hours, then moved to the lab bench in daylight for a further 24 hours. Germinated seeds were then planted in John Innes peat-based F2 compost in 40-well trays. Trays were placed in a Conviron controlled environment room with a 16-hour photoperiod: day temperature 18°C, night temperature 12°C, photosynthetic photon flux density (PPFD) of 350 µE/m^2^ at plant height. When the second leaf was fully expanded, usually at around 14 days after germination, inoculum was prepared.

Sporulating cultures of *Z. tritici* were grown on potato dextrose agar (PDA) plates for five to seven days under near ultra-violet light (Snijders Micro Clima-Series(tm) Economic Lux Chamber, Snijders Labs, Tilburg, The Netherlands) for 16 h per day at 18°C. Cultures were then flooded with 3 ml of sterile distilled water and scraped to release conidia. The concentration of conidial suspension was then adjusted to the desired inoculum concentration, typically 10^6^ spores ml^−1^. Conidial concentration was assessed through the use of a Fuchs-Rosenthal counting chamber (Hawksley, Lancing, UK). Two drops of polyoxyethylene-sorbitan monolaurate (Tween-20; Sigma-Aldrich Chemie Gmbh, Germany) were added per 50 ml of spore suspension.

Later-formed leaves were cut away so that only the primary seedling leaf remained. Seedlings were then evenly sprayed with spore suspension (20 ml per tray), assisted by the use of a turn table (made at JIC), using a Clarke Wiz Mini Air Compressor spray gun kit (Clarke Tools, Dunstable, England).

Trays were placed on matting within propagators, with two trays per propagator. This allowed trays to be watered from underneath to prevent the inoculum from washing off. The propagators were closed and covered with a black plastic bag for dark incubation. Black bags were removed after 48 hours and propagator lids were kept over trays until seven days after inoculation to increase humidity and therefore the success of infection by *Z. tritici*. New leaf growth was cut back twice per week to keep the inoculated leaves healthy and facilitate scoring.

The percentage of leaf area covered by pycnidia and leaf damage was scored by eye four to six times at intervals of two to five days over a period of 10 to 32 days post inoculation, depending on disease progress. Damage was defined as the combined area of necrosis and chlorosis. For imaging, leaves were mounted on A4 paper and scanned with a Canon LiDE 120 scanner at 600 DPI using the Canon IJ Scan Utility2 software. The standardised A4 image size allowed the dimensions of cropped images to be calculated using Adobe Illustrator.

### Statistical analysis

The area under the disease progress curve (AUDPC) was calculated for each dataset by calculating the area of the trapezium formed between each pair of scoring days on a graph of disease severity over time. The data were analysed for the effects of line, isolate and experimental design factors using linear mixed modelling to account for both random and fixed effects, via the package lmerTest^54^ in R version 4.2.2. If only fixed effects were involved, the native R analysis of variance aov() function was used. Nested deviance tests were conducted to determine the most concise fixed models that explained as much of the variation in phenotype as possible. The quality of models was assessed by residual plots. Models were fitted to the percentage of the maximum possible AUDPC but if the residual plots indicated non-normality or heteroscedasticity, AUDPC was transformed by the empirical logit transformation using the smallest possible AUDPC value score as a coefficient to avoid logarithms of zero^55,56^. Generally, it was possible to analyse damage data on the original percentage scale, whereas pynicidial coverage usually required logit transformation. The estimated mean pycnidia and damage scores for each genotype were obtained through the R emmeans^57^ package. These calculations were performed in R^58^ version 4.2.2.

### Genome-wide association study from Watkins collection

The markers used for GWAS of Watkins collection were ∼10 Mb core SNPs generated from whole genome shotgun sequencing of accessions and alignment to Chinese Spring. Extreme outlier values of phenotypic data were removed. In addition, we calculate kinship matrix as the covariate using GEMMA-kin. Based on these, we performed GWAS using GEMMA (v0.98.1) with parameters (gemma-0.98.1-linux-static -miss 0.9 -gk kinship.txt) and gemma- 0.98.1-linux-static -miss 0.9 -lmm -k kinship.txt). In-house scripts programmed in R were used to visualize these results.

### Estimation of haplotypes/alleles of candidate genes

A python script was written to identify the haplotypes of the six candidate *Stb* genes in the 6AS locus. The script parsed variant call format files (VCFs) generated from the alignment of Watkins and wheat lines to Chinese Spring (see above). This produced a matrix of distances between all accessions which could be used to determine haplotype groups. The R package pheatmap^59^ was used to generate heatmaps arranged in dendrograms from distance matrices, including associated phenotype data. Various iterations of the VCF parsing script described above were run and plotted in order to identify the most useful variation for haplotype calling. Ultimately, the whole gene sequence was analysed (rather than, for example, exons alone). The dendrogram produced was manually analysed to estimate the number of haplotype groups present. Clusters were then estimated using the cutree function in pheatmap and examined; several iterations were performed to determine the number of clusters/haplotypes which were most informative, particularly for explaining phenotypes. This method was used to determine which Watkins landraces carry the functional Arina allele of the *Stb15* candidate as well as the functional Chinese Spring allele of *Stb6*.

### *Generation of figures presenting* Stb *gene alleles in the Watkins collection*

Figures wherein *Stb* gene alleles were plotted (1a, 3a-b) were generated in R using ggplot2^60^ and cowplot^61^. For Figure 3a, the R package ggmap^62^ was implemented for generating the map and plotting coordinates.

A principal components analysis (PCA) was conducted in R version 4.2.2 using the package vcfR^63^ to process the WatSeq VCF data for *Stb15*, the base R prcomp function to compute the PCA and the vegan^64^ package for further analysis.

Composite main figures and illustrations were generated in Adobe Illustrator 2023.

### *Identification of candidate* Stb *genes by bioinformatics*

Candidate Septoria resistance genes were identified by selecting the most likely candidate from the genes in the LD block most highly associated with Septoria response. A number of factors were considered, such as: the SNP *p*-value (for association with Septoria response), gene class, the presence of differential SNPs between susceptible and resistant wheat varieties, and the strength of correlation of predicted resistant haplotypes with STB responses (described above).

To further confirm the *Stb15* candidate bioinformatically, a second iteration of the GWAS was run with lines carrying predicted functional alleles of the candidate gene removed. This resulted in the loss of the association of the 6AS locus with resistance, implying that the lines that were removed did contain the 6AS resistance.

### Generation of an Arina EMS population

Generation of the Arina EMS population is as described in Kolodziej et al. (2021)^48^. EMS mutagenesis of cv. Arina was performed with a concentration of 0.6% and 0.45% EMS (Sigma Aldrich, St. Louis, Missouri, USA), respectively. Seeds were incubated for 16 h in water at 4 °C, dried for 8 h on filter paper, and incubated for 16 h with shaking at room temperature in EMS solution. After washing three times for 30, 45, and 60 min, respectively, and for another 30 min under running tap water, seeds were pre-germinated on humid filter paper. Three thousand seeds of BC2F5-85 were mutagenized and pre-germinated seeds were propagated in the field. Single spikes of M0 plants were harvested and M1 plants were grown and harvested in the field.

### *Validation of the* Stb15 *candidate through screening an Arina EMS population*

When available, 12 seeds per M_2_ family were sown in a mixture of 1/2 blond and 1/2 brown peat mosses (Humustar soil, NPK 14-16-18, SARL Activert, Riom, France) and kept at 6°C for 4 days. Subsequently, the plants were cultivated in a growth chamber equipped with sodium lamps (HQI-TS 250W/D UVS FC2 FLH1, intensity = 300 µmol m^−2^ s^−1^) under a photoperiod of 16 h of light, a temperature of 21°C/18°C (day/night), and a relative humidity of 85%. Fourteen days after sowing, the plants were inoculated by spraying them with a hand sprayer (Elyte 2, Berthoud) with *Z. tritici* isolate IPO88004. The plants were covered with plastic bags for 3 days before returning to normal conditions. Visual evaluations were conducted at 21 and 28 days post-inoculation (dpi). All M_2_ plants carrying pycnidia were self-pollinated. M_3_ plants were evaluated for resistance to isolate IPO88004, following the procedure described above, with the exception that inoculations were performed on six-centimeter sections in the middle of the second leaf using a paintbrush. Three plants per M_3_ family were inoculated during two independent experiments. The *Z. tritici* inoculum was prepared using YG and YPD media following the procedure described in Battache et al.^28^. Inoculation with concentrations of 1 × 10^6^ spores/ml and 1 × 10^7^ spores/ml, supplemented with 0.05% (v/v) Tween-20 were used for inoculating M_2_ and M_3_ plants, respectively. The *Stb15* candidate gene was sequenced from each susceptible M_3_ plant using Sanger sequencing following PCR amplification using primer pairs Stb15F1/Stb15R1 and Stb15F3/Stb15R3 and the Phusion High-Fidelity Master Mix.

## Primers

Stb15F1: TCCTACTACTAGCCAAGCATGTC

Stb15R1: GCCATTGCCGTTAGAAACAG

Stb15F3: CTGTTCGAGGGAGGTTCCTA

Stb15R3: GTGCAAAGACCGCAGTATGT

### *Design of the* Stb15 *binary vector construct*

A wheat transformation vector was assembled using standard Golden Gate MoClo assembly (Werner et al., 2012) and traditional digestion and ligation cloning. The level 1 plasmids pL1P1R *Pv*UbiP:*hpt*-int:35sT selection cassett, pICH47742 L1P2 MCS & LacZ (Addgene #48001), and pL1P3ZmUbiP:GRF-GIF:NosT (Addgene #198047) were assembled into the Level 2 acceptor pGoldenGreenGate-M (pGGG-M) (Addgene #165422) binary vector (Smedley et al., 2021) along with end linker pELE-3 (Addgene #48018). The resulting plasmid was deemed pGGG L2 PvUH GGLacZ GRF-GIF. The *Stb15* gene sequence was analysed using the software Geneious Prime version 2020.2.4 (Biomatters) and two restriction enzymes (*Sbf*I and *Sac*I) were chosen for digestion/ligation cloning. The sequence containing the *Stb*15 gene (6077 bp), consisting of 1,917 bp promoter, 136 bp 5’UTR, 2,290 bp CDS, 306 bp 3’UTR and 1,404 bp terminator, was synthesised (Invitrogen, Thermofisher Scientific) with restriction enzyme recognition sites *Sbf*I and *Sac*I added to the 5’ and 3’ ends, respectively. The *Stb*15 gene synthon was cloned into pGGG L2 PvUH GGLacZ GRF-GIF within the multiple cloning site (MCS) using *Sbf*I and *Sac*I digestion/ligation. The resulting plasmid was named pGGG L2 *TaStb*15 and was electroporated into the hypervirulent *Agrobacterium* strain AGL1 (Lazo et al.,1991) as previously describe by Hayta et al., (2019).

### Agrobacterium *transformation of* T. aestivum *cv. ‘Fielder’*

Wheat transformation was performed as previously published by Hayta et al., (2021) with slight modification. The construct incorporated the GRF4-GIF1 technology (Debernardi et al., 2020). Briefly, wheat cv. ‘Fielder’ was grown in a controlled environmental room under a long-day photoperiod (16 h at 600 µmol m^−2^ s^−1^ light, at 20 °C day and 16 °C night). Wheat spikes were collected ∼14 days post anthesis (early milk stage GS73) when the immature embryos were 1-1.5 mm in diameter. Under aseptic conditions, immature embryos were isolated from surface sterilised grain.

The isolated immature embryos were pre-treated by centrifugation in liquid medium prior to *Agrobacterium* inoculation. The embryos were transferred to co-cultivation medium, scutellum side up, and incubated at 24 °C in the dark for 3 days co-cultivation. The embryogenic axes were excised and discarded, before transferring the embryos to wheat callus induction (WCI) medium without selection for 5 days at 24 °C in the dark. After 5 days, the embryos were transferred to WCI containing 15 mg L^−1^ hygromycin and incubated at 24 °C in the dark. Subculturing onto fresh WCI with hygromycin selection at 15 mg L^−1^ occurred every two weeks over a 5-week period. For the final, 5^th^, week on WCI the cultures were maintained in low light conditions at 24 °C. The cultures were transferred onto wheat regeneration medium (WRM) supplemented with 0.5 mg L^−1^ zeatin, and 15 mg L^−1^ hygromycin in deep petri dishes (90 mm diameter × 20 mm) and cultured under full fluorescent lights (100 µM m^−2^ s^−1^) with a 16 h photoperiod. Regenerated plantlets were transferred to De Wit culture tubes (Duchefa-Biochemie, W1607) containing rooting medium supplemented with 20 mg L^−1^ hygromycin. After approximately 10 days, rooted plants were transferred to soil (John Innes cereal mix in 24CT trays) and acclimatised (Hayta et al., 2019). The transgenic plants were maintained under the same growing conditions as donor material with a long-day photoperiod (16 h at 600 µmol m^−2^ s^−1^ light, at 20 °C day and 16 °C night). Transgenesis was confirmed and transgene copy number analysis performed using Taqman qPCR and probe as described in Hayta et al., (2019). The values obtained were used to calculate transgene copy number according to published methods (Livak and Schmittgen, 2001).

### *Experimental validation of the* Stb15 *transgenics*

The experiment to test the effect of the *Stb15* candidate on Septoria leaf blotch symptoms caused by the *Stb15*-avirulent *Z. tritici* isolate IPO88004 was sown in five 40-well seed trays with two experimental replicates per tray in a randomised design (ten replicates per line). Conditions and infection protocol were as described for JIC pathology experiments above. The percentage leaf area covered by lesions containing pycnidia was analysed by linear mixed modelling of repeated measures. Pycnidial area was logit-transformed to reduce heteroscedasticity. An antedependence order 1 model of repeated measures of logit (pycnidial area) at 21, 25, 29 and 32 days post inoculation (DPI) was used with the individual plant as the experimental subject. The random effect model was Tray (Replicate within Tray was omitted from the model because it did not cause a significant amount of random variation) and the fixed effects model was DPI * (Gene / Line), where the factor Gene indicates whether a wheat line contains *Stb15* either as the candidate transgene or by breeding, or alternatively lacks *Stb15*. Statistical analysis was done with Genstat 22nd edition (VSN International, Hemel Hempstead, UK).

### *Identification of intron-exon structure of* Stb15

RNAseq data for Arina*LrFor* generated by Anthony Hall and Manuel Spannagl was mapped to Arina*LrFor* using BWA^65^ version 0.7.7 and further processed and indexed via samtools^66^ version 1.2. The alignment was then visualised and intron/exon structure was manually annotated in IGV version 2.14.0. This was then confirmed by comparison with annotations generated by Anthony Hall and Manuel Spannagl based on the same RNAseq data.

### *Protein structure prediction of* Stb15

The protein structures encoded by the Stb15 candidate genes in Chinese Spring and Arina were predicted using AlphaFold (version 2.2.0)^67^. For each protein sequence, the highest-confidence prediction was selected for further analysis. Each protein sequence was also annotated using InterProScan^68^. These annotations were visualised on each protein’s structures using PyMol (version 2.5.2) and domain boundaries were manually expanded upon to include unannotated amino acid.

### KASP genotyping of European cultivars

KASP genotyping was carried out as described in Saintenac et al. (2021)^4^ on 278 European wheat cultivars. The marker sequences were as follows:

F = GAAGGTGACCAAGTTCATGCTGGTTTCAACTTGCAATATGATC

V = GAAGGTCGGAGTCAACGGATTGGTTTCAACTTGCCATATGATT

C = AGTGAACCAGGTGCCAAAAC

### Analyses of sequence evolution

#### Identification of potential Stb15 homologs in plants

High-quality reference genome protein annotations of plant species were downloaded (**Supp. Table 9**). Local blastp databases were generated using command line BLAST^69^ version 2.13.0. The amino acid sequence of the functional Stb15 allele from Arina*LrFor* was used as a query sequence for BLAST searches against protein databases of each species. The top 30 hits were recovered to ensure that no potential orthologs or paralogs were missed.

#### Protein alignment

A protein alignment was generated using MUSCLE^70^ version 3.8.31 with default settings. Sequences were removed if they contained large (>350 aa) and divergent insertions which disrupted the alignment or if they contained less than two of the domains present in the Arina*LrFor* Stb15 sequence. Multiple splice variants were included if their predicted amino acid sequence varied.

#### *Phylogenetic tree construction and analysis of the* Stb15 *clade*

ModelFinder^71^ was used to predict the best evolutionary model for the alignment (JTT+R10) implemented via IQ-TREE^72^ version 1.6.10. Branch supports were obtained with ultra-fast bootstrap (UFBoot2^73^) and tree reconstruction was performed using IQ-TREE. The least-repetitive clade containing *Stb15* was extracted and sequence conservation was analysed in Geneious version 2022.2.2. Genome annotations (GFFs) for each ortholog were used to draw gene structures in R version 4.2.2. For the purpose of the figures presented (**Fig. 3c** and **Supp. Fig. 5**), all exon annotations for each gene are presented within a single leaf and splice variants were pruned from the tree. The tree image was generated using iTOL^74^ version 6.

#### *Alignment for consensus sequence of inner* Stb15 *clade*

A small alignment of the inner *Stb15* clade was generated by MUSCLE within Geneious version 2022.2.2. It was noticed that the *Zea mays* homolog Zea_mays_Zm00001eb119590_P002 was in fact a tandem duplication containing two identical sequences of a protein encoding a partial BTL and full SLG, PAN and S/TPK domains. To reduce disruption of the alignment, one half of this protein sequence was retained. A screenshot of the consensus chart from the Geneious alignment was used in **Fig. 3d**.

## References

1. Savary, S. et al. The global burden of pathogens and pests on major food crops. Nat Ecol Evol 3, 430–439 (2019).

2. O’Driscoll, A., Kildea, S., Doohan, F., Spink, J. & Mullins, E. The wheat–Septoria conflict: a new front opening up? Trends Plant Sci 19, 602–610 (2014).

3. Saintenac, C. et al. Wheat receptor-kinase-like protein Stb6 controls gene-for-gene resistance to fungal pathogen Zymoseptoria tritici. Nat Genet 50, 368–374 (2018).

4. Saintenac, C. et al. A wheat cysteine-rich receptor-like kinase confersbroad-spectrum resistance against Septoria tritici blotch. 6 (2021).

5. Salamini, F., Özkan, H., Brandolini, A., Schäfer-Pregl, R. & Martin, W. Genetics and geography of wild cereal domestication in the near east. Nat Rev Genet 3, 429–441 (2002).

6. Stukenbrock, E. H., Banke, S., Javan-Nikkhah, M. & McDonald, B. A. Origin and domestication of the fungal wheat pathogen Mycosphaerella graminicola via sympatric speciation. Mol Biol Evol 24, 398–411 (2007).

7. Hafeez, A. N. et al. Creation and judicious application of a wheat resistance gene atlas. Molecular Plant vol. 14 1053–1070 Preprint at 10.1016/j.molp.2021.05.014 (2021).

8. Jones, J. D. G. & Dangl, J. L. The plant immune system. Nature vol. 444 323–329 Preprint at 10.1038/nature05286 (2006).

9. Stotz, H. U., Mitrousia, G. K., de Wit, P. J. G. M. & Fitt, B. D. L. Effector-triggered defence against apoplastic fungal pathogens. Trends in Plant Science vol. 19 491–500 Preprint at 10.1016/j.tplants.2014.04.009 (2014).

10. Cook, D. E., Mesarich, C. H. & Thomma, B. P. H. J. Understanding Plant Immunity as a Surveillance System to Detect Invasion. Annu Rev Phytopathol 53, 541–563 (2015).

11. Kanyuka, K. & Rudd, J. J. Cell surface immune receptors: the guardians of the plant’s extracellular spaces. Curr Opin Plant Biol 50, 1–8 (2019).

12. Kema, G. H. J., Yu, D., Rijkenberg, F. H. J., Shaw, M. W. & Baayen, R. P. Histology of pathogenesis of Mycosphaerella graminicola in wheat. Phytopathology vol. 7 777–786 Preprint at 10.1094/Phyto-86-777 (1996).

13. Shaw, M. W. Assessment of upward movement of rain splash using a fluorescent tracer method and its application to the epidemiology of cereal pathogens. Plant Pathol 36, 201–213 (1987).

14. Brown, J. K. M., Chartrain, L., Lasserre-Zuber, P. & Saintenac, C. Genetics of resistance to Zymoseptoria tritici and applications to wheat breeding. Fungal Genetics and Biology vol. 79 33–41 Preprint at 10.1016/j.fgb.2015.04.017 (2015).

15. Yang, N., Mcdonald, M. C., Peter Solomon, S. & Milgate, A. W. Genetic mapping of Stb19, a new resistance gene to Zymoseptoria tritici in wheat. Theoretical and Applied Genetics 1, 3 (2018).

16. Langlands-Perry, C. et al. Resistance of the Wheat Cultivar ‘Renan’ to Septoria Leaf Blotch Explained by a Combination of Strain Specific and Strain Non-Specific QTL Mapped on an Ultra-Dense Genetic Map. Genes (Basel) 13, 100 (2022).

17. Brading, P. A., Verstappen, E. C. P., Kema, G. H. J. & Brown, J. K. M. A Gene-for-Gene Relationship Between Wheat and Mycosphaerella graminicola, the Septoria Tritici Blotch Pathogen. Phytopathology 92, 439–445 (2002).

18. Ghaffary, S. M. T. et al. New broad-spectrum resistance to septoria tritici blotch derived from synthetic hexaploid wheat. Theoretical and Applied Genetics 124, 125–142 (2012).

19. Arraiano, L. S. et al. A gene in European wheat cultivars for resistance to an African isolate of Mycosphaerella graminicola. Plant Pathol 56, 73–78 (2007).

20. Arraiano, L. S. & Brown, J. K. M. Identification of isolate-specific and partial resistance to septoria tritici blotch in 238 European wheat cultivars and breeding lines. Plant Pathol 55, 726–738 (2006).

21. Wingen, L. U. et al. Establishing the A. E. Watkins landrace cultivar collection as a resource for systematic gene discovery in bread wheat. Theor Appl Genet 127, 1831–1842 (2014).

22. Winfield, M. O. et al. High-density genotyping of the A.E. Watkins Collection of hexaploid landraces identifies a large molecular diversity compared to elite bread wheat. Plant Biotechnol J 16, 165–175 (2018).

23. Togninalli, M. et al. The AraGWAS Catalog: A curated and standardized Arabidopsis thaliana GWAS catalog. Nucleic Acids Res 46, D1150–D1156 (2018).

24. Arora, S. et al. A wheat kinase and immune receptor form host-specificity barriers against the blast fungus. Nat Plants 9, (2023).

25. Kema, G. H. J. et al. Variation for virulence and resistance in the wheat-Mycosphaerella graminicola pathosystem I. Interactions between pathogen isolates and host cultivars. Phytopathology vol. 86 213–220 Preprint at 10.1094/Phyto-86-213 (1996).

26. Chartrain, L., Sourdille, P., Bernard, M. & Brown, J. K. M. Identification and location of Stb9, a gene for resistance to septoria tritici blotch in wheat cultivars Courtot and Tonic. Plant Pathol 58, 547–555 (2009).

27. Amezrou, R. et al. A secreted protease-like protein in Zymoseptoria tritici is responsible for avirulence on Stb9 resistance gene in wheat. PLoS Pathog 19, (2023).

28. Battache, M. et al. Blocked at the Stomatal Gate, a Key Step of Wheat Stb16q-Mediated Resistance to Zymoseptoria tritici. Front Plant Sci 13, (2022).

29. Sun, Y., Qiao, Z., Muchero, W. & Chen, J. G. Lectin Receptor-Like Kinases: The Sensor and Mediator at the Plant Cell Surface. Front Plant Sci 11, 1989 (2020).

30. Avni, R. et al. Genome sequences of three Aegilops species of the section Sitopsis reveal phylogenetic relationships and provide resources for wheat improvement. Plant J 110, 179–192 (2022).

31. Bartoli, C. & Roux, F. Genome-Wide Association Studies In Plant Pathosystems: Toward an Ecological Genomics Approach. Front Plant Sci 8, 763 (2017).

32. Chartrain, L., Brading, P. A. & Brown, J. K. M. Presence of the Stb6 gene for resistance to septoria tritici blotch (Mycosphaerella graminicola) in cultivars used in wheat-breeding programmes worldwide. Plant Pathol 54, 134–143 (2005).

33. Steuernagel, B. et al. The NLR-Annotator Tool Enables Annotation of the Intracellular Immune Receptor Repertoire 1[OPEN]. (2020) doi:10.1104/pp.19.01273.

34. Wulff, B. B. H., Chakrabarti, A. & Jones, D. A. Recognitional specificity and evolution in the tomato-Cladosporium fulvum pathosystem. Molecular Plant-Microbe Interactions 22, 1191–1202 (2009).

35. Thomas, C. M., Dixon, M. S., Parniske, M., Golstein, C. & Jones, J. D. G. Genetic and molecular analysis of tomato Cf genes for resistance to Cladosporium fulvum. Philosophical Transactions of the Royal Society B: Biological Sciences 353, 1413–1424 (1998).

36. Larkan, N. J. et al. The Brassica napus wall-associated kinase-like (WAKL) gene Rlm9 provides race-specific blackleg resistance. The Plant Journal 104, 892–900 (2020).

37. Larkan, N. J. et al. The Brassica napus blackleg resistance gene LepR3 encodes a receptor-like protein triggered by the Leptosphaeria maculans effector AVRLM1. New Phytologist 197, 595–605 (2013).

38. Miyakawa, T. et al. A secreted protein with plant-specific cysteine-rich motif functions as a mannose-binding lectin that exhibits antifungal activity. Plant Physiol 166, 766–778 (2014).

39. Burton, R. A. & Fincher, G. B. Evolution and development of cell walls in cereal grains. Front Plant Sci 5, 456 (2014).

40. Singh, P. & Zimmerli, L. Lectin receptor kinases in plant innate immunity. Front Plant Sci 4, 124 (2013).

41. Wang, Y. et al. Orthologous receptor kinases quantitatively affect the host status of barley to leaf rust fungi. Nat Plants 5, 1129–1135 (2019).

42. van der Hoorn, R. A. L. & Kamoun, S. From Guard to Decoy: A New Model for Perception of Plant Pathogen Effectors. THE PLANT CELL ONLINE 20, 2009–2017 (2008).

43. Ma, L. S. et al. Maize Antifungal Protein AFP1 Elevates Fungal Chitin Levels by Targeting Chitin Deacetylases and Other Glycoproteins. mBio 14, (2023).

44. Bouwmeester, K. et al. The Lectin Receptor Kinase LecRK-I.9 Is a Novel Phytophthora Resistance Component and a Potential Host Target for a RXLR Effector. PLoS Pathog 7, e1001327 (2011).

45. Amezrou, R. et al. Whole-genome sequencing reveals diverse mechanisms underlying quantitative pathogenicity and host adaptation in a fungal plant pathogen. bioRxiv 2022.12.23.521735 (2022) doi:10.1101/2022.12.23.521735.

46. Zhong, Z. et al. A small secreted protein in Zymoseptoria tritici is responsible for avirulence on wheat cultivars carrying the Stb6 resistance gene. New Phytologist 214, 619–631 (2017).

47. Kema, G. H. J. et al. Stress and sexual reproduction affect the dynamics of the wheat pathogen effector AvrStb6 and strobilurin resistance. Nat Genet 1 (2018) doi:10.1038/s41588-018-0052-9.

48. Kolodziej, M. C. et al. A membrane-bound ankyrin repeat protein confers race-specific leaf rust disease resistance in wheat. Nature Communications 2021 12:1 12, 1–12 (2021).

49. Kema, G. H. J. & Van Silfhout, C. H. Genetic variation for virulence and resistance in the wheat-Mycosphaerella graminicola pathosystem III.Comparative seedling and adult plant experiments. Phytopathology 87, 266–272 (1997).

50. Kema, G. H. J. et al. Genetic variation for virulence and resistance in the wheat-Mycosphaerella graminicola pathosystem I. Interactions Between Pathogen Isolates and Host Cultivars. Phytopathology 86, 200–212 (1996).

51. Kema, G. H., Sayoud, R., Annone, J. G. & Van Silfhout, C. H. Genetic Variation for Virulence and Resistance in the Wheat-Mycosphaerella graminicola Pathosystem II. Analysis of Interactions Between Pathogen Isolates and Host Cultivars. Phytopathology vol. 86 213–220 Preprint at (1996).

52. Patterson, H. D. & Williams, E. R. A new class of resolvable incomplete block designs. Biometrika 63, 83–92 (1976).

53. Arraiano, L. S., Brading, P. A. & Brown, J. K. M. A detached seedling leaf technique to study resistance to Mycosphaerella graminicola (anamorph Septoria tritici) in wheat. Plant Pathol 50, 339–346 (2001).

54. Kuznetsova, A., Brockhoff, P. B. & Christensen, R. H. B. lmerTest Package: Tests in Linear Mixed Effects Models. J Stat Softw 82, (2017).

55. McGrann, G. R. D. et al. A trade off between mlo resistance to powdery mildew and increased susceptibility of barley to a newly important disease, Ramularia leaf spot. J Exp Bot 65, 1025–1037 (2014).

56. Collett, D. Modelling binary data BT - Texts in statistical science. (Chapman & Hall/CRC, 2003).

57. Lenth, Russel., Love, Jonathon. & Herve, Maxime. Estimated Marginal Means, aka Least-Squares Means. R J (2018).

58. R Core Team. R: A language and environment for statistical computing. International journal of antimicrobial agents vol. 41 Preprint at <https://www.R-project.org/ (2022).

59. Kolde, R. Package pheatmap’. Bioconductor (2012).

60. Wickham, H. Package ggplot2: Elegant Graphics for Data Analysis. Springer-Verlag New York (2016).

61. Wilke, C. O. cowplot: Streamlined Plot Theme and Plot Annotations for ‘ggplot2’. R package version 0.7.0. URL: https://CRAN.R-project.org/package=cowplot. xAcessed 18 October 2018 (2016).

62. Kahle, D. & Wickham, H. ggmap: Spatial Visualization with ggplot2. https://journal.r-project.org/archive/2013-1/kahle-wickham.pdf.

63. Knaus, B. J. & Grünwald, N. J. vcfr: a package to manipulate and visualize variant call format data in R. in Molecular Ecology Resources vol. 17 (2017).

64. Oksanen, J. et al. Vegan: Community Ecology Package. R package version 2.5-7. Community ecology package 10, (2020).

65. Li, H. Aligning sequence reads, clone sequences and assembly contigs with BWA-MEM. arXiv preprint arXiv (2013).

66. Danecek, P. et al. Twelve years of SAMtools and BCFtools. Gigascience 10, (2021).

67. Jumper, J. et al. Highly accurate protein structure prediction with AlphaFold. Nature 596, (2021).

68. Paysan-Lafosse, T. et al. InterPro in 2022. Nucleic Acids Res 51, (2023).

69. Altschul, S. F., Gish, W., Miller, W., Myers, E. W. & Lipman, D. J. Basic local alignment search tool. J Mol Biol 215, (1990).

70. Edgar, R. C. MUSCLE: Multiple sequence alignment with high accuracy and high throughput. Nucleic Acids Res 32, (2004).

71. Kalyaanamoorthy, S., Minh, B. Q., Wong, T. K. F., Von Haeseler, A. & Jermiin, L. S. ModelFinder: fast model selection for accurate phylogenetic estimates. Nature Methods 2017 14:6 14, 587–589 (2017).

72. Nguyen, L. T., Schmidt, H. A., Von Haeseler, A. & Minh, B. Q. IQ-TREE: A Fast and Effective Stochastic Algorithm for Estimating Maximum-Likelihood Phylogenies. Mol Biol Evol 32, 268–274 (2015).

73. Hoang, D. T., Chernomor, O., Von Haeseler, A., Minh, B. Q. & Vinh, L. S. UFBoot2: Improving the Ultrafast Bootstrap Approximation. Mol Biol Evol 35, 518–522 (2018).

74. Letunic, I. & Bork, P. Interactive Tree Of Life (iTOL) v5: an online tool for phylogenetic tree display and annotation. Nucleic Acids Res 49, W293–W296 (2021).

75. International Wheat Genome Sequencing Consortium (IWGSC), T. I. W. G. S. C. et al. Shifting the limits in wheat research and breeding using a fully annotated reference genome. Science 361, eaar7191 (2018).

