## Supplementary Figures, Tables and Text for "Septoria tritici blotch resistance gene *Stb15* encodes a lectin receptor-like kinase"

### Stb15 paper supplementary information

### Supplementary Figures

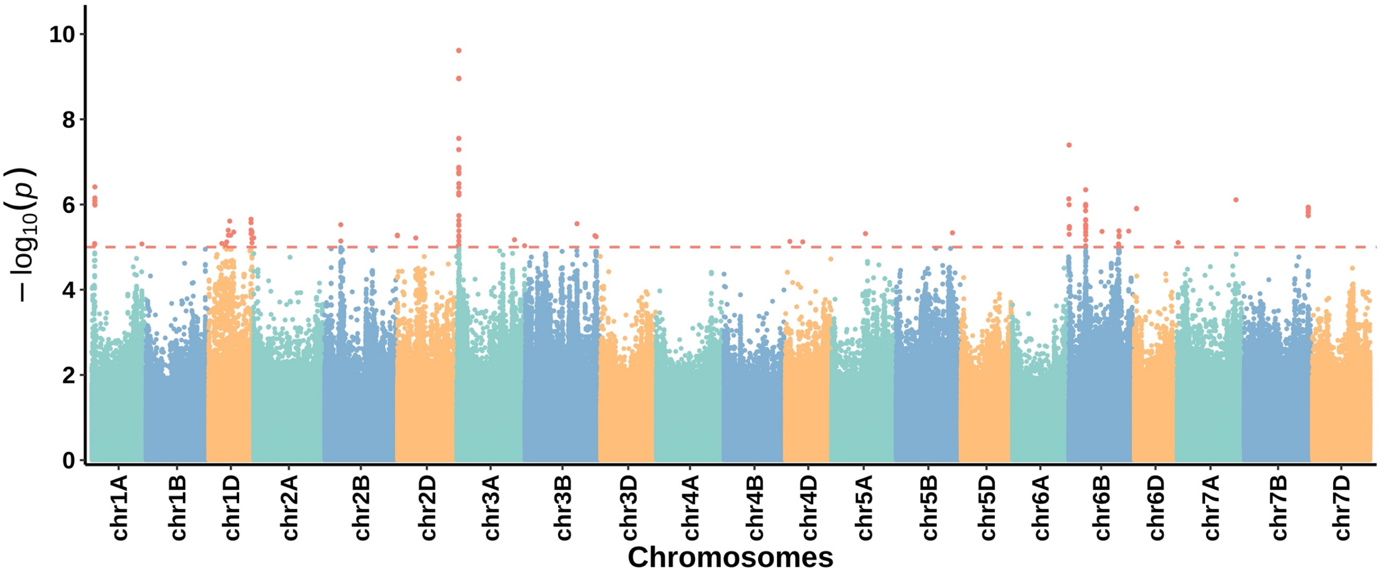


Supplementary Fig. 1: Manhattan plot showing associations between SNPs in the Watkins landrace collection and damage (% maximum dAUDPC) in response to *Z. tritici* isolate IPO323.

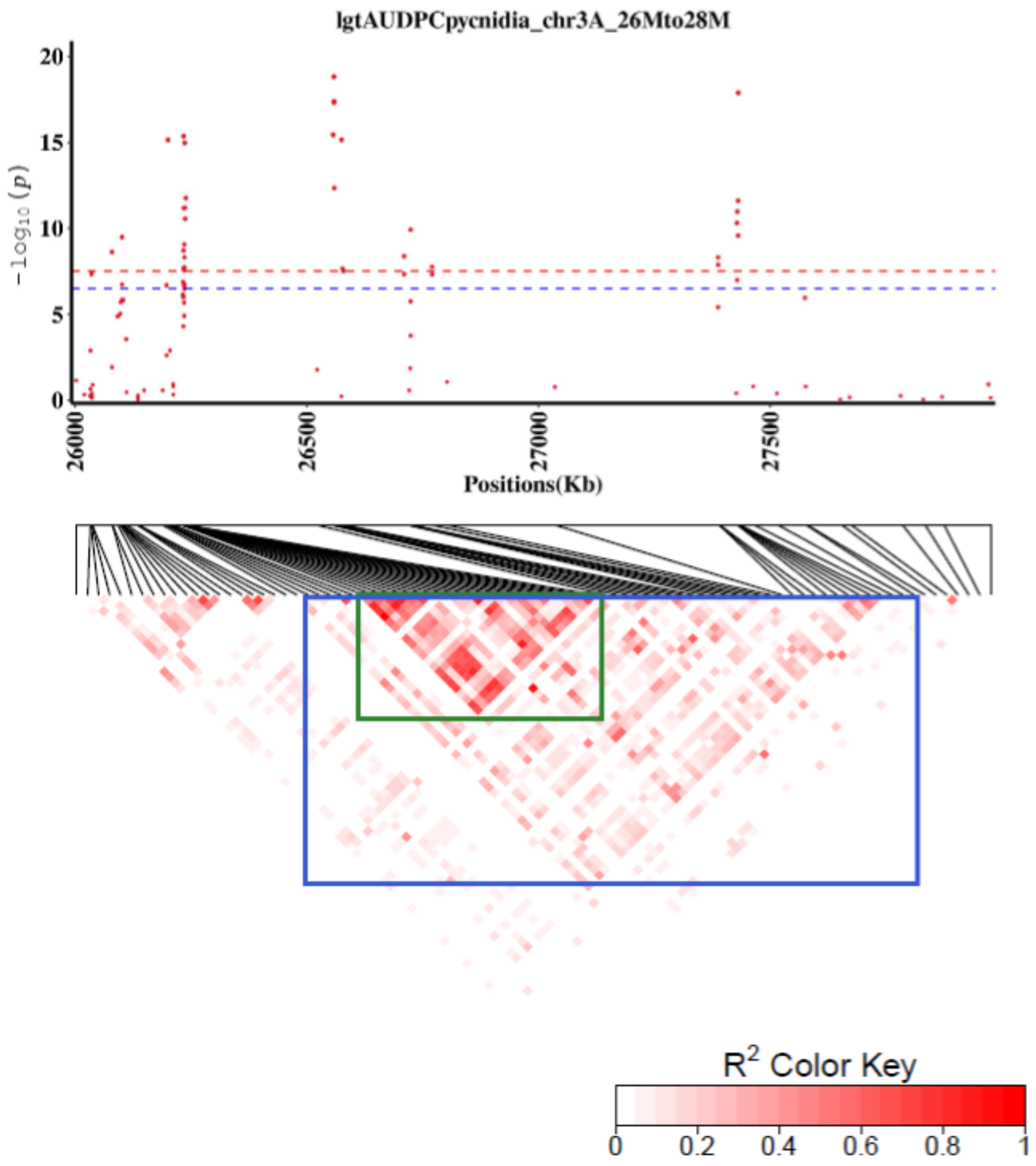


**Supplementary Fig. 2:** Zoomed-in Manhattan plot of the 3A peak from the SNP-based GWAS with the IPO323 pycnidia phenotype. The linkage disequilibrium heatmap is displayed below. Blue square from 26.10 to 27.50 Mb indicates the main area of the peak. Green square from 26,035,170 to 26,238,727 bp indicates the smaller haploblock within the blue square which is most highly associated with resistance.

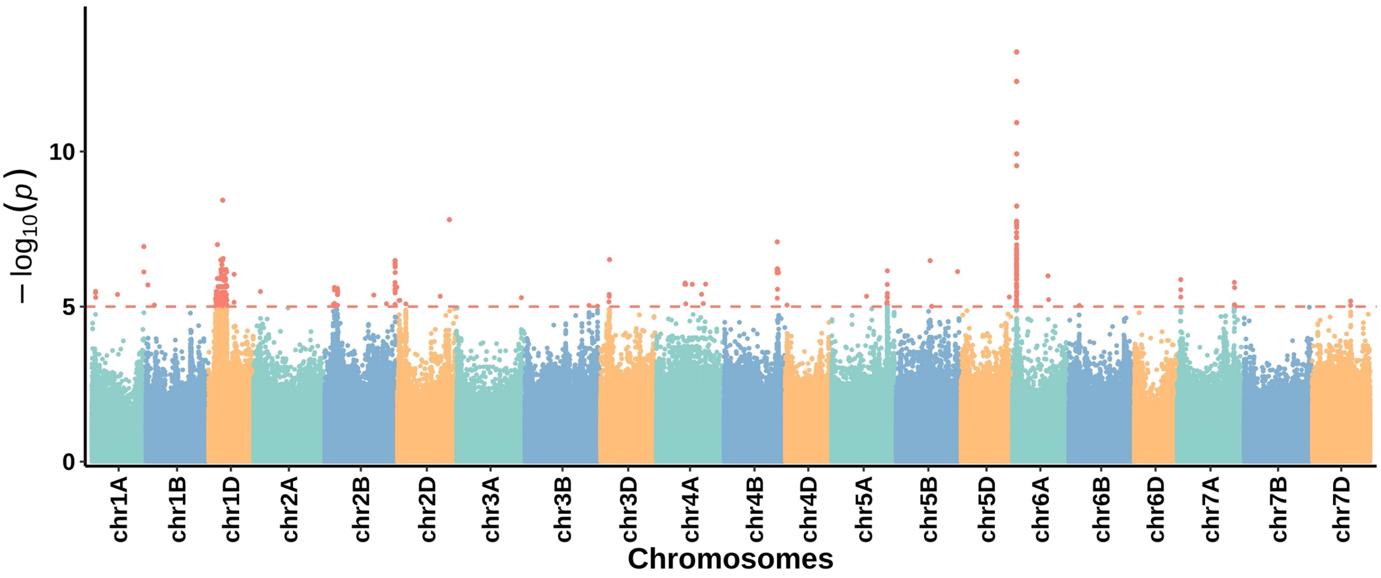


Supplementary Fig. 3: Manhattan plot showing association of SNPs mapped to Chinese Spring with damage (% maximum dAUDPC) in response to *Z. tritici* isolate IPO88004.

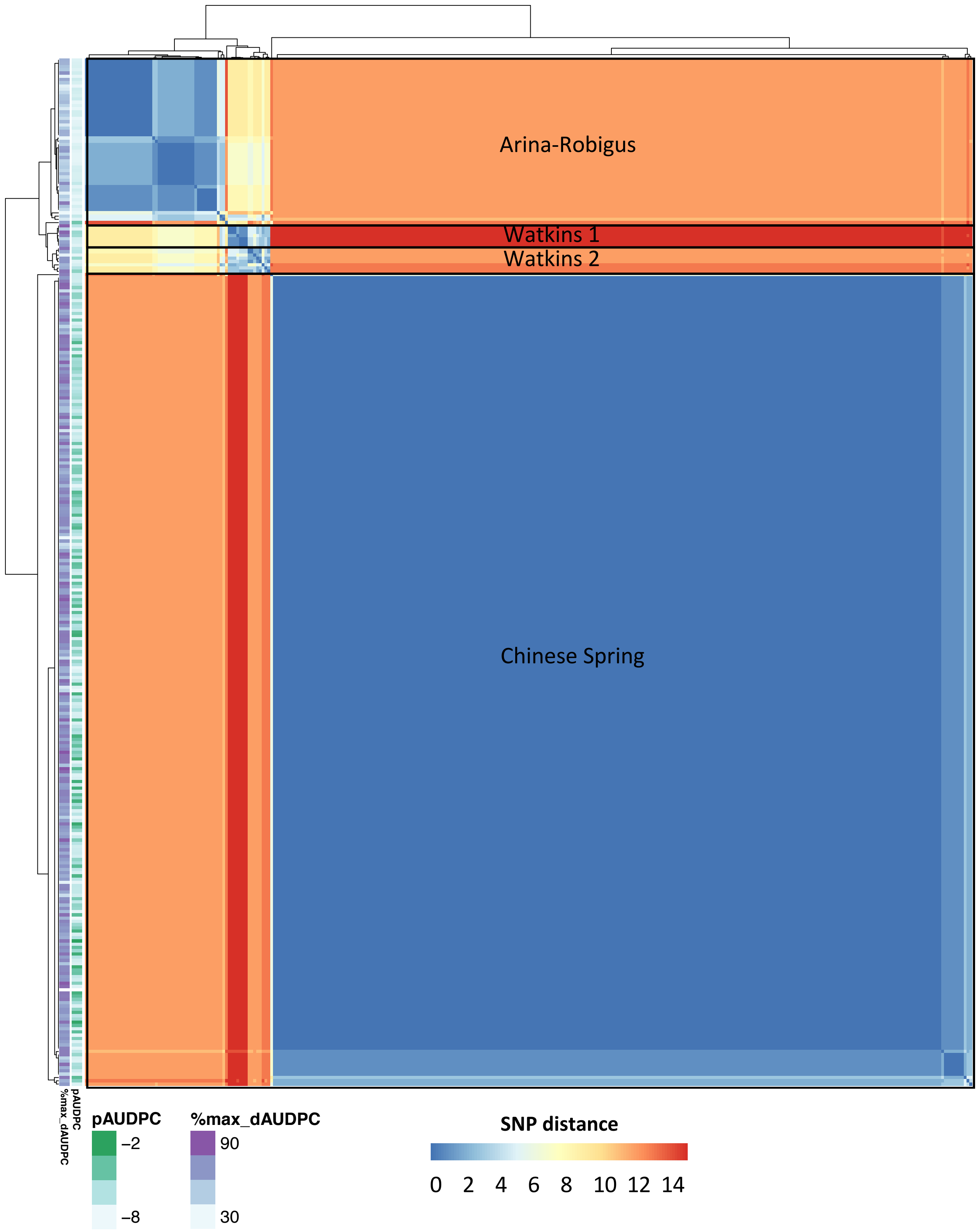


**Supplementary Fig, 4**: A matrix of the distance in SNPs between Watkins accessions within the TraesCS6A02G078700 locus rendered as a heat map. SNPs were called across a panel of 300 Watkins accessions in relation to landrace Chinese Spring (WatSeq consortium) and SNP distance calculated using a custom programme. Panel on the left shows phenotype scores: logit pAUDPC (pycnidia) in green and % max. dAUDPC (damage) in purple. Major haplotype groups are outlined in black and labelled with the names of key lines within those groups.

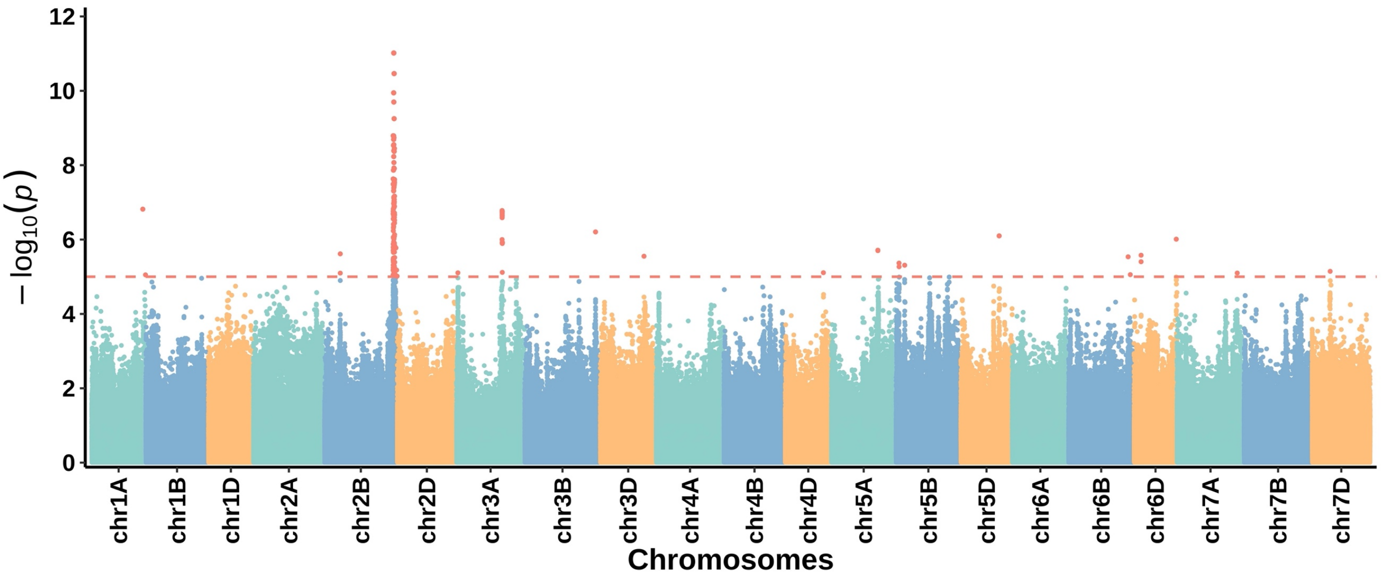


**Supplementary Fig 5**: Manhattan plot of Watkins SNPs associated with IPO88004 pycnidia data, with lines carrying the functional allele of *Stb15* candidate TraesCS6A02G078700 removed.

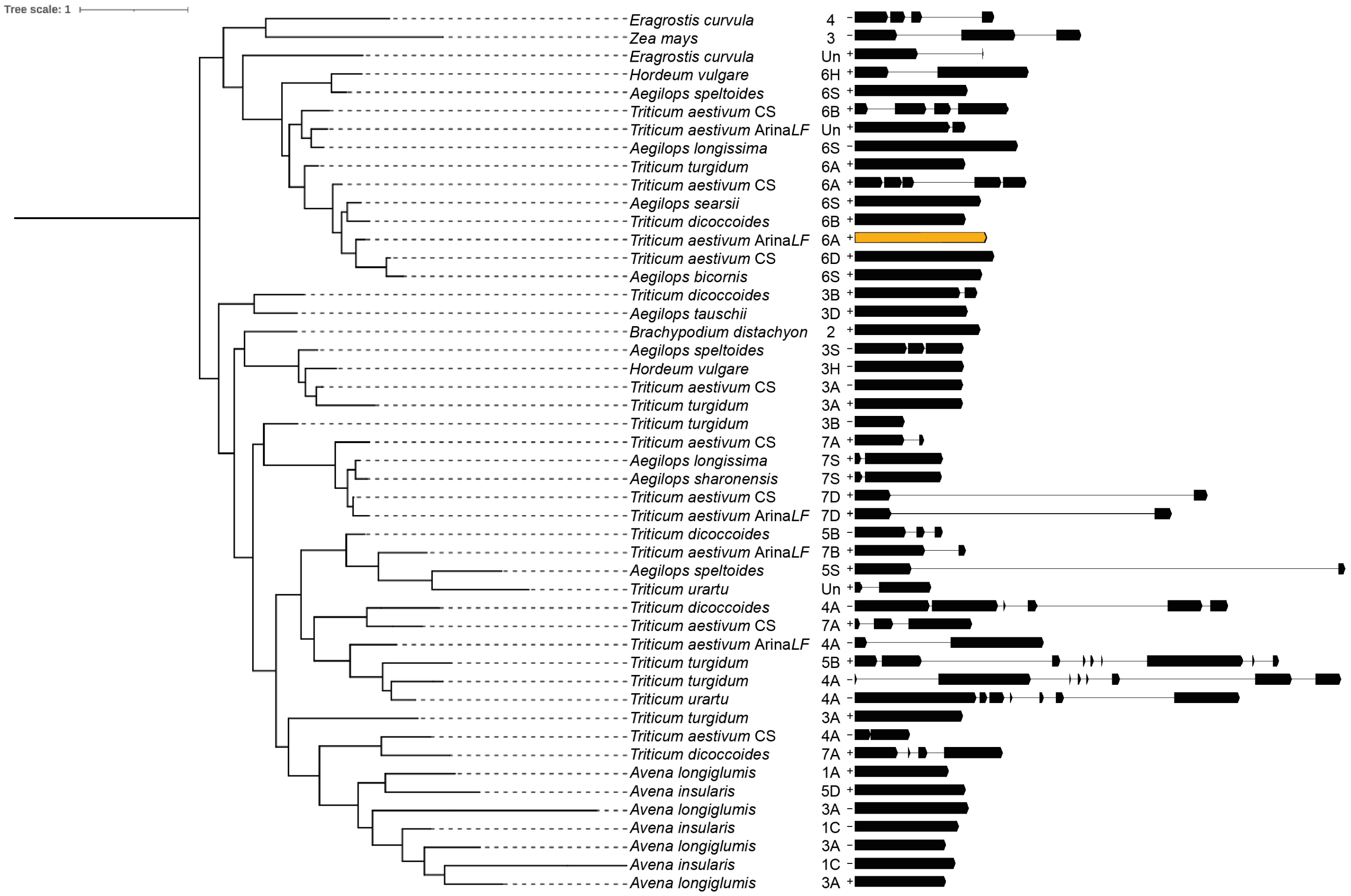


**Supplementary Fig. 6**: Phylogenetic tree clade containing *Stb15* generated from a dataset of the top 30 BLAST protein hits to the Arina*LrFor* allele of *Stb15* (denoted Arina*LF* in tree) from 32 plant species with high-quality published genome sequences. This clade features 16 Poaceae species. Gene structures are given on the right based on annotation files for each genome. Arrows indicate exons whilst lines indicate introns. *Stb15* is highlighted in orange.

### Supplementary Tables

**Supplementary Table 1:** ANOVA table of linear mixed model for logit pAUDPC scores from the Watkins 300 collection inoculated with IPO323, IPO88004 and IPO90012. Colons represent nested factors.

| Term | Mean Square | Numerator DF | Denominator DF | F value | Pr(>F) |
|---|---|---|---|---|---|
| Isolate | 21.45 | 2.00 | 3.10 | 8.53 | 0.05 |
| Line | 28.49 | 322.00 | 3539.20 | 11.34 | <0.0001 |
| Scorer | 5.89 | 2.00 | 46.80 | 2.34 | 0.11 |
| Isolate:Line | 14.40 | 624.00 | 3535.20 | 5.73 | <0.0001 |

**Supplementary Table 2:** ANOVA table of linear mixed model for % maximum dAUDPC scores from the Watkins 300 collection inoculated with IPO323, IPO88004 and IPO90012. Colons represent nested factors.

| Term | Mean Square | Numerator DF | Denominator DF | F value | Pr(>F) |
|---|---|---|---|---|---|
| Isolate | 607.48 | 2.00 | 3.00 | 12.46 | 0.03 |
| Line | 427.76 | 322.00 | 3523.50 | 8.77 | <0.0001 |
| Scorer | 3.78 | 2.00 | 60.10 | 0.08 | 0.93 |
| Isolate:Line | 219.63 | 624.00 | 3502.20 | 4.51 | <0.0001 |

**Supplementary Table 3:** Random effects of the linear mixed model fitted to logit pAUDPC and % maximum dAUDPC scores form the Watkins 300 collection inoculated with IPO323, IPO88004 and IPO90012. Colons represent nested factors.

| | Logit pAUDPC | | % max. dAUDPC | |
|---|---|---|---|---|
| Term | Variance | Std. Dev | Variance | Std. Dev |
| Isolate:Batch:Rep:Box:Tray | | | 0.54 | 0.74 |
| Isolate:Batch:Rep:Box | 0.11 | 0.33 | 2.19 | 1.48 |
| Isolate:Batch:Rep | 0.01 | 0.12 | 0.00 | 0.00 |
| Isolate:Batch | 0.37 | 0.61 | 23.56 | 4.85 |
| Residual | 2.51 | 1.59 | 48.75 | 6.98 |

**Supplementary Table 4:** see SupplementaryTables.xlx**.** Table of estimated logit pAUDPC and percentage of the maximum dAUDPC scores of the Watkins and wheat lines tested against IPO323, IPO88004 and IPO90012. Mean response of lines to all isolates is provided. Some wheat controls were only included in two assays so do not have a line mean estimate across all isolates, but a mean calculated from mean responses to the tested isolates is given in italics (this was not estimated via linear mixed modelling). Haplotypes of *Stb6* and *Stb15* inferred from WatSeq SNPs are also provided. For *Stb6*, haplotypes 1 and 2 are known resistant haplotypes whilst for *Stb15* A/R is known to be resistant and CS is susceptible.

**Supplementary Table 5:** Wheat lines included in each Septoria assay and reasons for their inclusion (yes = included). The top section of the table includes wheat lines whose genomes have been sequenced by IWGSC (International Wheat Genome Sequencing Consortium (IWGSC) et al., 2018) or as part of the wheat pangenome project (Walkowiak et al., 2020). Below are selections based on information from Arraiano & Brown (2006), Brown et al. (2015) and Chartrain et al. (2004). The bottom panel consists of cv. Fielder and results from this project.

| Line | IPO323 | IPO88004 | IPO90012 | Reason for inclusion |
|---|---|---|---|---|
| Chinese Spring | Yes | Yes | Yes | Wheat reference genome |
| Arina*LrFor* | Yes | Yes | Yes | Wheat pangenome |
| Baj | Yes | Yes | Yes | Wheat cultivar |
| Cadenza | Yes | Yes | Yes | Wheat pangenome |
| CDC Landmark | Yes | Yes | Yes | Wheat pangenome |
| CDC Stanley | No | Yes | Yes | Wheat pangenome |
| Claire | No | Yes | Yes | Wheat pangenome |
| Kronos | No | Yes | Yes | Wheat pangenome |
| Jagger | No | Yes | Yes | Wheat pangenome |
| Julius | No | Yes | Yes | Wheat pangenome |
| Lancer | Yes | Yes | Yes | Wheat pangenome |
| Mace | No | Yes | Yes | Wheat pangenome |
| Norin 61 | No | Yes | Yes | Wheat pangenome |
| Paragon | Yes | Yes | Yes | Wheat pangenome |
| Robigus | Yes | Yes | Yes | Wheat pangenome |
| SY Mattis | No | Yes | Yes | Wheat pangenome |
| Weebill | No | Yes | Yes | Wheat pangenome |
| Courtot | No | No | Yes | Susceptible to IPO90012 |
| Flame | Yes | No | No | Resistant to IPO323 |
| Longbow | Yes | Yes | Yes | Widely susceptible |
| Olaf | No | No | Yes | Resistant to IPO90012 |
| Fielder | No | Yes | No | Susceptible to IPO88004, background of transgenics |

**Supplementary Table 6:** Genes within the region on chromosome 6A associated with resistance to *Z. tritici* isolate IPO88004 and their function. Reasons for not selecting these genes as the *Stb15* candidate are given.

| Annotation | Gene Name | Start | Length (bp) | Protein | Reason for exclusion |
|---|---|---|---|---|---|
| **TRAESCS6A02G078600** | APUM23 | 48516680 | 5780 | Pumilio homolog 23 | Arina*LrFor* has the same genotype as Chinese Spring. |
| **TRAESCS6A02G079000** | S6PDH | 48563464 | 3903 | Aldo-ket-red domain-containing protein | Association of haplotypes with resistance phenotypes is not strong. |
| **TRAESCS6A02G078800** | PEX16 | 48552372 | 4331 | Peroxisomal membrane protein PEX16 | No SNPs in exons detected. |
| **TRAESCS6A02G078900** | SRK6 | 48556992 | 4149 | Uncharacterised protein | Association of haplotypes with resistance phenotypes is not strong. |
| **TRAESCS6A02G078700** | | 48525265 | 3354 | Receptor-like serine/threonine-protein kinase | **No reason to exclude** – clear association of the Arina*LrFor* haplotype group with resistance (**Supplementary Figure 4**). |
| **TRAESCS6A02G078500** | | 48509308 | 1890 | Uncharacterised, LRR superfamily related domain | Arina*LrFor* has the same genotype as Chinese Spring. |

Supplementary Table 7: Pycnidia scores from Arraiano et al. (2009) that demonstrates that resistance to IPO88004 does not correlate with IPO89011, and therefore that the 2BL resistance to IPO88004 is distinct from *Stb9* resistance to IPO89011. Emboldening and asterisks indicate specific resistance.

| Line | IPO88004 | IPO89011 | Conclusion |
|---|---|---|---|
| Sportsman | **1*** | 14 | Resistant to IPO88004 but not to IPO89011 |
| Selkirk | **0*** | 21 | |
| Tipstaff | **1*** | 35 | |
| Carstens VIII | **0*** | 36 | |
| Maris Settler | **0*** | 30 | |
| Courtout | 67 | **12*** | Resistant to IPO89011 but not to IPO88004 |
| Melbor | 27 | **1*** | |
| Tonic | 20 | **5*** | |
| Jena | 47 | **1*** | |
| Soissons | 45 | **6*** | |
| Plus 59 more lines | | | |

**Supplementary Table 8:** REML variance components analysis of fixed terms for the repeated measures analysis conducted on Fielder transgenic $T_2$ lines containing *Stb15*, nulls and wheat varieties inoculated with *Z. tritici* isolate IPO88004. The fixed effect model specified was DPI * (Gene / Line) whilst the random model was Tray.

| Term | Numerator DF | Denominator DF | F value | P value |
|---|---|---|---|---|
| DPI | 1 | 157.0 | 21.87 | <0.001 |
| Gene | 5 | 103.9 | 9.25 | <0.001 |
| DPI:Gene | 5 | 157.0 | 1.26 | 0.284 |
| Gene:Line | 11 | 104.2 | 2.59 | 0.006 |
| DPI:Gene:Line | 11 | 157.0 | 1.08 | 0.378 |

**Supplementary Table 9**: Table of means for Fielder transgenic $T_2$ lines containing *Stb15,* nulls and wheat varieties inoculated with *Z. tritici* isolate IPO88004 estimated from the model presented in **Supplementary Table 8**. Lines are presented in order with most resistant lines at the top and most susceptible at the bottom. Transgenic lines have a number code of X.Y which refers to the $T_0$ family (X) and $T_1$ parent (Y). CNX:Y refers to the copy number of $T_0$ (X) and $T_1$ (Y) parents. Copy number of $T_2$ plants is estimated from $T_1$ parents.

| Line | $T_0$ copy number | $T_2$ copy number (est) | Estimated Mean | SE |
|---|---|---|---|---|
| 21.1 CN4:7 | 4 | 6-8 | -6.2598759 | 0.50557824 |
| Arina | - | - | -6.0761606 | 0.39431071 |
| BastardII | - | - | -6.0421435 | 0.37333641 |
| 22.6 CN2:4 | 2 | 4 | -6.0381205 | 0.35599329 |
| 06.9 CN1:2 | 1 | 2 | -5.9841042 | 0.37332816 |
| Longbow | - | - | -5.9063111 | 0.34122667 |
| 21.8 CN4:2 | 4 | 0-4 | -5.7521766 | 0.37486134 |
| 25.10 CN1:2 | 1 | 2 | -5.712293 | 0.35592845 |
| 25.1 CN1:2 | 1 | 2 | -5.423515 | 0.42215545 |
| 21.2 CN4:5 | 4 | 2-8 | -5.3373171 | 0.45629993 |
| 25.7 CN1:0 | 1 | 0 | -5.1949022 | 0.35592845 |
| 01.6 CN2-GRF:4 | 2 (construct only) | 4 (construct only) | -5.1646468 | 0.42410934 |
| 11.10 CN2-GRF:4 | 2 (construct only) | 4 (construct only) | -5.0494157 | 0.42417409 |
| Fielder | - | - | -4.9830822 | 0.39430519 |
| Chinese Spring | - | - | -4.2276224 | 0.39575282 |
| 06.5 CN1:0 | 1 | 0 | -3.9315759 | 0.35599329 |
| 30.7 CN0 | 0 | 0 | -3.877265 | 0.39430519 |

Supplementary Table 10: SNPs identified from the WatSeq alignment against Chinese Spring within *Stb6*.

| Genomic position | Chinese Spring allele | Alternative allele |
|---|---|---|
| 26,200,567 | T | C |
| 26,200,618 | C | G |
| 26,200,653 | A | G |
| 26,200,659 | G | A |
| 26,201,022 | A | `G |
| 26,201,037 | C | T |
| 26,201,046 | T | C |
| 26,201,734 | G | A |
| 26,202,969 | G | A |
| 26,203,022 | A | G |
| 26,203,076 | A | T |
| 26,203,115 | A | G |
| 26,203,148 | C | A |
| 26,203,155 | G | T |
| 26,203,192 | G | T |
| 26,203,263 | C | T |
| 26,203,264 | A | G |
| 26,203,430 | C | T |
| 26,205,300 | A | T |
| 26,205,344 | T | C |
| 26,206,172 | T | A |
| 26,206,287 | T | G |

**Supplementary Table 11:** SNPs identified from the WatSeq alignment against Chinese Spring within the TraesCS6A02G078700 (*Stb15*) locus. SNP genotype for each haplotype defined from **Supplementary Figure 6** is given. Robigus has the same genotype as Arina/Arina*LrFor* at this locus (based on wheat pangenome assemblies) but due to some areas of low read coverage there were missing datapoints in this locus for Arina*LrFor*, resulting in differences in SNP distance between Robigus and Arina*LrFor*. The more complete SNP dataset for Robigus can therefore be referred to here as the reference for the functional resistant allele of *Stb15.*

| | | Distance from | | SNP position chr6A_part1:4852… | | | | | | | | | | | | | | |
|---|---|---|---|---|---|---|---|---|---|---|---|---|---|---|---|---|---|---|
| | | | | 1 | 2 | 3 | 4 | 5 | 6 | 7 | 8 | 9 | 10 | 11 | 12 | 13 | 14 | 15 |
| **Haplotype** | **No.** | **Chinese Spring** | **Arina*LrFor*** | 5467 | 5473 | 5514 | 5805 | 5907 | 5922 | 5923 | 5925 | 6158 | 6193 | 7920 | 8004 | 8102 | 8609 | 8693 |
| Chinese Spring | 250 | 0-6 | 11-12 | C | C | C | A | C | C | T | A | C | A | G | G | C | C | T |
| Robigus | 51 | 11-12 | 0-7 | A | G | C | A | A | A | C | G | T | C | A | T | A | C | C |
| Watkins 1 | 7 | 15 | 11 | A | G | C | . | . | . | . | . | . | . | A | T | A | T | C |
| Watkins 2 | 7 | 12 | 10-11 | A | G | T | G | C | C | T | G | . | . | A | T | A | T | C |
| | | | | EXON 1 | | | | EXON 2 | | | | | | EXON 4 | | | EXON 5 | |

**Supplementary Table 12:** KASP genotyping of *Stb15* in European wheat cultivars. See SupplementaryTables.xlsx.
**Supplementary Table 13:** Genomes used for evolutionary analyses. See SupplementaryTables.xlsx file.
**Supplementary Table 14**: Protein and gene IDs of *Stb15* homologs in **Fig. 3c**. See SupplementaryTables.xlsx file.

### Supplementary Text

Supplementary Text 1: Country abbreviations used in Figure 3.

AFG = Afghanistan, ALB = Albania, ALG = Algeria, ARM = Armenia, AUS = Australia, AZE = Azerbaijan, B&H = Bosnia & Herzegovina, BUL = Bulgaria, CHI = China, CI = Canary Islands, CRO = Croatia, CYP = Cyprus, EGY = Egypt, ETH = Ethiopia, FRA = France, FIN = Finland, GEO = Georgia, GRE = Greece, HUN = Hungary, IND = India, IRN = Iran, IRQ = Iraq, ITA = Italy, KAZ = Kazakhstan, LEB = Lebanon, MOR = Morocco, MYA = Myanmar, NM = North Macedonia, PAK = Pakistan, PAL = Palestine, POL = Poland, POR = Portugal, ROM = Romania, RUS = Russia, SER = Serbia, SPA = Spain, SYR = Syria, TKM = Turkmenistan, TUN = Tunisia, TUR = Turkey, UKR = Ukraine, UK = United Kingdom.
